# Within-Family GWAS does not Ameliorate the Decline in Prediction Accuracy across Populations

**DOI:** 10.1101/2024.12.12.628188

**Authors:** Luyin Zhang, Dalton Conley

**Affiliations:** Office of Population Research; Princeton University; New York Genome Center, New York, NY; National Bureau of Economic Research, Cambridge MA

## Abstract

As polygenic prediction extends beyond the research domain to involve clinical applications, the urgency of solving the “portability problem” becomes amplified—that is, the fact that polygenic indices (PGI) constructed based on discovery analysis in one population (typically of exclusively continental European descent) predict poorly in other populations. In the present paper we test whether population differences in genetic nurture, assortative mating, or population stratification contribute to the fact that polygenic indices constructed based on GWAS results from European-descent samples predict more poorly in admixed populations with Native American and African ancestry. We do this by comparing the rates of decline in prediction accuracy of classical-GWAS-based PGIs versus within-family-based PGIs, each estimated in a population of European descent, as they are deployed in two samples of Latino Americans and African Americans. Within-family GWAS putatively eliminates the effects of parental genetic nurture, assortative mating, and population stratification; thus, we can determine whether without those confounding factors in the PGI construction, the relative prediction accuracy in the out-groups is ameliorated. Results show that relative prediction accuracy is not improved, suggesting that the differences across groups can be almost entirely explained by variation in genetic architecture (i.e. allele frequencies and short-range LD) rather than the aforementioned factors. Additional analysis of the impact of genetic architecture on the decline in prediction accuracy supports this conclusion. Future researchers should test within-family analysis at the prediction rather than the discovery stage.

## Introduction

Since the first polygenic risk score was calculated in 2009 for the phenotype of schizophrenia (International Schizophrenia Consortium 2009), the deployment of these summative measures of the effects of common genetic variants has grown exponentially. In the years since that first genome-wide risk score paper was published, over twenty-five thousand have been published using the terms *polygenic risk score* (*PRS*), *genome-wide risk score* (*GRS*), *polygenic score* (*PGS*), or *polygenic index* (PGI – our preferred term in this paper). The deployment of this tool shows no signs of abating in the human genetics research world. Moreover, novel clinical applications of polygenic indices are emerging, from prevention (i.e. prescribing statins for individuals with high scores for cardiovascular disease long before hypercholesterolemia or other symptoms appear [O’Sullivan et al. 2022]) to diagnosis (i.e. distinguishing depression or ADHD subtypes through PGI profiling [Ruderfer et al. 2018]) to embryo selection in fertility medicine (i.e. optimizing the known genetic potential for certain traits or risks [Turley et al. 2021; Kumar et al. 2022]). More use cases are surely on the horizon.

Given the rapidly growing utilization of PGIs, it is all the more troubling that the vast majority of PGIs have been constructed based on genome-wide association study (GWAS) weights of samples composed of individuals of exclusively European continental descent (Martin et al. 2020). Since European countries and countries populated by a majority of people of European descent (e.g., the United States, Canada, Australia) tend to enjoy adequate resources to genotype large numbers of people in surveys, biobanks, direct-to-consumer genetic testing, and other data collection modes, this is, perhaps, understandable. Recently, East Asian countries have been accelerating the pace of genetic data collection (Mills and Rahal 2019; Petersen et al. 2019), but other ancestry groups remain far behind in terms of representation in GWAS. For example, only two percent of all genetic data represent populations of sub-Saharan African ancestry, even though these populations represent 16 percent of the total world population and, in fact, display the most genetic diversity of any continental group (United Nations 2024).

The underrepresentation of certain genetic ancestries is problematic given the fact that extensive research has shown that the polygenic indices constructed based on discovery analysis of populations of exclusively European ancestry lose prediction accuracy when applied to other genetic-similarity groups (Martin et al. 2017, 2020). This “portability problem” is most acute for prediction within populations of African descent (*ibid*), with Native American populations exhibiting the second most inaccurate prediction. For example, European-based PGIs predict populations of African descent to be systematically shorter than people of European descent, a shift so dramatic that the two (predicted) distributions barely overlap (Martin et al. 2017), even though this does not match the empirical reality of observed phenotypes in the world. Recent research has shown that prediction accuracy declines even within broadly genetically similar groups as a function of genetic distance (Ding et al. 2023) or even as a function of social position within a sample of individuals of similar genetic relatedness (Mostafavi et al. 2020)

There has been much discussion of the roots of this portability problem. There are a number of dynamics that may contribute to the phenomenon. These include issues of genetic architecture such as allele frequency differences across groups and inter-population variation in linkage disequilibrium that leads to differential association of measured SNPs with causal alleles. But they also include the possibility that confounds baked into classical GWAS analysis—deriving from gene-environment correlation, long-range LD induced by assortative mating, and genetic nurture—may be operating differently across the spectrum of genetic similarity.

Some recent work has used admixed subjects (African Americans) and shown that the prediction accuracy of European-based PGIs is a function of the proportion European in the individual’s genome (Bitarello and Mathieson 2020). This is highly suggestive evidence that it is genetic architecture and not environmental confounds that explains the deterioration of prediction accuracy when PGIs are ported across populations since. However, those with more (or less) African (or European) genetic similarity proportions probably experience systematically differing environments (and rates of genetic assortative mating), even when their race is shared, as evidenced by, for example, a robust literature on the “pigmentocracy” in the African American community (see, e.g., Monk Jr. 2021).

In the present paper, we build on Bitarello and Mathieson (2020) by comparing the decline in prediction accuracy for PGIs based on GWAS weights calculated from analysis of European-descent populations when applied to people with Native American admixture (Latino Americans) and when applied to people with sub-Saharan African ancestry (African Americans). We compare the relative prediction accuracy of PGIs calculated with classical (i.e. between-family) GWASs and from family-based (i.e. within-family) GWASs (Tan et al. 2024). The latter GWAS is meant to factor out population stratification (Price et al. 2006), the genetic signature of assortative mating (Beauchamp 2016), and genetic nurture (Kong et al. 2018), leaving only the estimate of the direct genetic effect. In this way, we can see if the relative decline in prediction accuracy is stemmed at all by using only the direct effects—which would suggest that factors like genetic nurture and gene-environment correlation may play a role in accounting for the portability problem.

We analyze five important phenotypes across two datasets. The phenotypes are height, body mass index, systolic blood pressure, education, and having ever smoked. The original Tan et al. (2024) GWAS on which we base our PGIs provided within-family and classical weights for 34 phenotypes; however, we were limited to those that appeared in both our test samples with sufficient sample size – the National Longitudinal Study of Adolescent to Adult Health (Add Health) and the Health and Retirement Study (HRS). This reduced the number of potential phenotypes to 11.^5^ When we performed the analysis, 6 of these phenotypes yielded problematic initial results—meaning that we could not come close to replicating the original decline in prediction accuracy in traditional models. Thus, we were left with the aforementioned five important phenotypes that range from health to physiological development to social and cognitive development. Most importantly, they range in the degree to which they display prediction accuracy decay when ported across populations and they vary in how much indirect genetic effects (i.e. population stratification, genetic nurture) contribute to the overall classical GWAS estimate when compared to the within-family estimate.

For example, among the five phenotypes under study, when ported from Europeans to African genetic similarity groups the decline in prediction accuracy of classical PGIs is 71.2 percent for height and 74.9 percent for BMI, whereas systolic blood pressure displays a diminution in prediction accuracy of 93.4 percent, ever smoked 91.8 percent, and education 82.3 percent. Meanwhile, in the Tan et al. (2024) analysis educational attainment shows the greatest “correction” in SNP heritability when within-family GWAS is substituted for classical, while BMI shows the least.

We chose Add Health and HRS because they are U.S.-based cohorts that each have respondents of multiple ancestries (and were not in the original Tan et al. GWAS). Add Health has the additional benefit of having a sibling-based subsample, which allows us to test the case where not just the discovery analysis involved within-family models but where the replication test is also performed within-family. (However, it turns out that the sample sizes for African Americans and Latinos in the sibling subsample are too low to generate any reasonably powered estimates, so we relegate those results to Figure S5 in the Online Supplemental Information).

To preview our results, we find that using within family GWAS weights, purged of many confounders that might vary across populations, does little to address the decline in prediction accuracy when ported across populations distinguished by their degree of genetic similarity. Further, we assess whether there is a pattern by which the relative prediction accuracy (for either PGI type, classically based or within-family) is associated with either assortative mating for that phenotype within both populations or the SNP heritability of the phenotype in both populations. We find that while portability does vary by these factors across this handful of phenotypes, the pattern is the same for classical or within-family based PGIs. Finally, we examine the extent to which the contribution of LD and allele frequency differences to the decline in prediction accuracy varies by the type of PGI. We find that the contribution of these measures of genetic architecture (for the two phenotypes for which we are able to estimate that contribution) does not vary by PGI type, further suggesting that it is allele frequency and short-range LD differences that are key to solving the problem of declining prediction accuracy across major population groups.

## Materials and Methods

### Data

#### Add Health

The National Longitudinal Study of Adolescent to Adult Health (Add Health) is a longitudinal study of a nationally representative sample of students from 132 middle and high school in the United States. The first wave of survey was fielded in 1994-1995 and consisted of 20,745 randomly selected students. This student cohort was followed with four additional in-home interviews in 1996, 2001-2002, 2008, and 2016-2018, where a rich set of sociodemographic, behavioral, psychosocial, familial, and contextual information was collected. In Wave 4, approximately 80% of the respondents who participated in the in-home survey (N = 15,701) provided saliva samples and consented to be genotyped. Two Illumina arrays – the Illumina Human Omni1-Quad BeadChip and the Illumina Human Omni-2.5 Quad BeadChip – were used for genotyping. After quality control procedures at the genetic marker and individual levels, Add Health kept 609,130 single nucleotide polymorphisms (SNPs) common to both chips for 9,974 respondents and imputed them to the 1000 Genomes (1KG) Phase 3 panel. The imputed genetic data are positioned in GRCh37 coordinates.

#### HRS

The Health and Retirement Study (HRS) is a biennial, longitudinal study initiated in 1992 that surveys a nationally representative sample of around 20,000 Americans over age 50 and their spouses. In 2006 (Phase 1), 2008 (Phase 2), and 2010 (Phase 3), saliva samples of consented participants were collected during enhanced face-to-face interviews conducted with a randomized subset of the HRS cohort. Genotyping was performed using the Illumina HumanOmni2.5-8v1 array. After standard quality controls, HRS collectively imputed genotype data from all three phases against the 1KG reference panel using IMPUTE2 (Howie et al. 2009). The imputed genetic data are in GRCh37 build and cover 15,620 HRS respondents, of which 15506 are unrelated.

#### SNP selection and phenotypic measures

Our primary analysis is restricted to bi-allelic SNPs that are of high imputation quality (R^2^ > 0.99), minor allele frequency (MAF) > 1%, a missing call rate < 5%, and in an ancestry-specific Hardy-Weinberg equilibrium (HWE) P < 10^-6^ (the HWE filtering step was performed after assignment of ancestral groups, a process detailed later). The stringent threshold for imputation quality was chosen to be consistent with Tan et al. (2024), our source of GWAS summary statistics, and to ensure the performance of subsequent imputation of parental genotypes for the Add Health siblings^6^ (Young et al. 2022).

We focus on five phenotypes that (1) are available in both datasets and (2) show reasonable prediction accuracy of PGI among our European subsamples. They include height, body mass index (BMI), systolic blood pressure (SBP), years of schooling, and having ever smoked. To minimize missingness on phenotypic measures, we used Wave 4 as our main source of information for the Add Health sample. Height was measured to the nearest 0.5 centimeters for all respondents who were capable of standing unassisted. BMI was calculated as weight measured to the nearest 0.1 kilograms divided by height in meters squared (BMI = kg/m2), after checking for inconsistencies between self-reported and measured weights or heights, sex, and waist. Add Health assessed SBP in millimeters of mercury through three serial measurements taken at 30-second intervals. The SBP value was constructed as the average of the second and third measures^7^ or the reading from the first one if both are unavailable. In cases where both measures were unavailable, the reading from the first measurement was employed. Following Tan et al. (2024), we mapped the self-reported highest educational attainment at Wave 4 to levels of the International Standard Classification of Education and subsequently converted them to the corresponding completed years of schooling. Ever smoked status was determined using answers to the following 2 questions: (1) “Have you ever smoked an entire cigarette?”, and (2) “Have you ever smoked cigarettes regularly, that is, at least 1 cigarette every day for 30 days?”

For the HRS sample, we primarily relied on the 1992-2020 longitudinal data pre-prepared by the RAND Center for the Study of Aging. Height was physically measured in inches during an enhanced face-to-face interview and converted to meters. BMI was derived as physically measured weight converted to kilograms divided by the square of height. Similar to Add Health, BP readings were taken three times during the in-home physical measurements portion of the HRS. The SBP value was constructed as the mean of the second and third measures when all three readings were valid and as the mean of all available readings otherwise. Years of education was determined by extracting the first non-missing value across all available waves and imputed with the information recorded in the HRS Tracker file if the former was absent. The ever-smoked question was typically only asked at the respondent’s first interview, and the answer was carried forward in subsequent waves unless a previous non-smoker indicated currently smoking.^8^

### Statistical Methods

#### Imputation of parental genotypes for Add Health

Leveraging sibling pairs from Add Health, we used the Mendelian imputation method implemented in snipar (Young et al. 2022) to infer the mean parental genotypes for each of them. We first aligned the quality-controlled SNPs to genomic build GRCh38 using LiftOver (Hinrichs et al. 2006) to match the built-in genetic map of snipar. This process led to a loss of 579 variants (N = 1,111,939) that could not be mapped. Full siblings were identified using KING (Manichaikul et al. 2010). We randomly kept one individual from each pair of monozygotic (MZ) twins if they have other non-MZ siblings genotyped. Otherwise, MZ twins were fully dropped as they do not provide enough genetic information for Mendelian imputation. After cross-checking the Add Health pedigree information with KING inferences, we identified 646 sibling pairs including 1,323 individuals. We used snipar to infer identity-by-descent (IBD) segments and subsequently impute parental genotypes. Since all input pairs are siblings (rather than parent-offspring), we can only obtain the average of maternal and paternal genotypes for each individual in this subsample.

#### Genetic ancestry inference & PC construction

We implemented principal component analysis (PCA) to identify genetic ancestral groups, largely following the practice in Yengo et al. (2018), and to create the first 10 principal component (PCs) for each group. Specifically, we ran first PCA on 2,490 unrelated individuals from the 1000 Genomes (1KG) Phase 3 that comprised 5 super populations: Africans (labeled as “AFR” in 1KG), Europeans (“EUR”), Admixed Americans (“AMR”), East Asians (“EAS”), and South Asians (“SAS”). Genetic variants used in the PCA were LD-pruned using a window size of 100kb, a step size of 5, and an R^2^ of 0.1 from an initial pool of bi-allelic HapMap 3 SNPs with MAF > 10% that were also available in our test samples (before ancestry-specific HWE filtering). After projecting our Add Health and HRS respondents onto the first two genetic PCs, we assigned them to the closest of the five 1KG reference populations based on Mahalanobis distance. Our subsequent analyses are limited only to European (EUR), American (AMR), and African (AFR) ancestral groups. Once these groups were identified, we implemented PCA again within each group to create the first 10 genetic PCs, which were later used in PGI analysis to adjust for possible population stratification.

#### Global ancestry proportion estimation

In addition to the categorical measure of genetic ancestral group described above, we also derived continuous measures of genetic ancestry that assigns an individual’s DNA proportionately to different genetic ancestral groups using ADMIXTURE (Alexander et al 2009). We call these measures genetic similarity proportions (GSP), in line with recent recommendations from the National Academies of Sciences, Engineering, and Medicine (2023). To run supervised ADMIXTURE, we retrieved unrelated individuals from the HapMap3 (International HapMap 3 Consortium 2010) and Human Genome Diversity Project (HGDP) (Bergström et al. 2020) as our reference panels for creating Sub-Saharan African (AFR), European (EUR), East Asian (EAS), and Indigenous American (IAM) GSPs. These includes 83 people from Yoruba in Nigeria (labeled as “YRI” in HapMap3), 36 Northern & Western Europeans in Utah (“CEU” in HapMap3), 137 Han Chinese in Beijing (“CHB” in HapMap3), and 34 people from Pima & Mayan in Mexico (“Pima” and “Maya” in HGDP). We limited ADMIXTURE analysis to quality-controlled genotyped autosomal SNPs that overlap between our test and reference samples and performed LD pruning in PLINK1.9 (Purcell et al. 2007) with a window size of 200kb, a step size of 25, and an R^2^ of 0.4.

The four GSPs obtained from supervised ADMIXTURE are highly consistent with estimates from unsupervised ADMIXTURE that specified four distinct ancestral populations to be modeled. Moreover, they highly align with results from Gnomix (Hilmarsson et al. 2021), a more computationally intensive approach for local ancestry estimation. We also cross-checked our genetic ancestral group assignment with the GSP measures by examining the distribution of the four GSPs within the EUR, AFR, and AMR group and found consistent and sensible patterns in both Add Health and HRS.

#### Heritability estimation

We used genetic data from the UK Biobank (UKB) to estimate the heritability of the five phenotypes for the European and African genetic ancestral groups. Following the strategy outlined above, we identified 452,172 and 7,993 people of European and African ancestry, respectively. To reduce computational time, heritability estimation for the European group was conducted on a random sample of 7,993 UKB respondents. For each ancestral group, we constructed genetic relationship matrices (GRMs) from autosomal HapMap3 SNPs with R^2^ > 0.99, MAF > 0.01, and ancestry-specific HWE P < 10^-6^ and removed cryptic relatedness using a cutoff value of 0.05. SNP heritability was then estimated with the GREML (Genomic Relatedness-Based Restricted Maximum Likelihood) method implemented in GCTA (Yang et al. 2011), controlling for age, sex, and the first 20 ancestry-specific genetic PCs.

#### PGI construction

We retrieved summary statistics for classical and family-based GWASs by Tan et al. (2024), which are publicly available in the Social Science Genetic Association Consortium. The summary statistics have been quality-controlled to include autosomal bi-allelic SNPs with high imputation quality (INFO score > 0.99) that has MAF > 1% and a call rate of >99%. We replicated our PGI construction and subsequent analyses using GWAS summary statistics from Howe et al. (2022), which included SNPs or INDELs with lower imputation quality (INFO score > 0.3) and adopted a within-sibship design. The findings are substantially similar to what we present below, based on Tan et al. (2024) and can be found in Figures S1-S4 of the Supplementary Material.

We used PRS-CS (Ge et al. 2019) to construct weights for polygenic scores (PGIs). PRS-CS is a high-dimensional Bayesian approach that utilizes a continuous shrinkage prior to account for LD and infer posterior SNP effects. We use the LD reference panel provided in PRS-CS, which comprises 375,120 individuals of European ancestry from the UKB and covers 1,117,425 HapMap3 SNPs. Global shrinkage parameters were learnt using a fully Bayesian approach. The obtained SNP weights were input to PLINK2 (Chang et al. 2015) for PGI calculation. We applied the same set of SNP weights to construct the mean parental PGIs for Add Health siblings using the pgs.py script of snipar (Young et al. 2022).

#### Prediction accuracy

Prediction accuracy was calculated as the square of partial correlation between the PGI and the corresponding phenotype (which is essentially the incremental R^2^ of the PGI) after they have been both residualized using sex, birth year, and the first 10 genetic PCs. For family-based PGI analysis using the Add Health sibling subsample, the covariates used for residualization further includes the mean parental PGI. In practice, we obtained the partial correlation and its standard error from regressing the standardized phenotype residuals on the standardized PGI residuals.

#### Phenotypic assortative mating

We calculated levels of assortative mating for the five phenotypes using spousal pairs from the HRS and siblings from Add Health, analyzing each dataset separately by genetic ancestry. In the HRS, we included only heterosexual spousal pairs from the same genetic ancestry group who were in their first marriage. This sample comprised 1,688 pairs of spouses of European ancestry and 193 pairs of African ancestry. Assortative mating in the HRS cohort was assessed by Pearson correlation. For Add Health, we estimated assortative mating using 805 full siblings of European ancestry and 224 of African descent. This was done by fitting random-effects models that nested individuals within families, controlling for birth year and gender, and calculating the intra-class correlation coefficient.

#### Loss of prediction accuracy due to LD and MAF differences

We used the method developed in Wang et al. 2020 to quantify the contribution of LD and MAF differences to the relative prediction accuracy of the PGI in Africans as compared to Europeans. We first estimated the predicted relative accuracy associated with LD and MAF differences *RA_pred(LD+MAF)_* from 1KG^9^ based on the equation (2) in Wang et al. (2020). We then derived the proportion of the loss of accuracy (LOA) explained by LD and MAF as 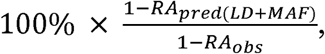 where *RA_obs_* refers to the AFR-to-EUR ratio of prediction accuracy obtained from our test samples.

## Results

In the Panel B of Figure 1, below, we show the prediction accuracy of classical and family-based PGIs among Latino and African American samples relative to the European sample for the five phenotypes. These results are meta-analyzed across two datasets, Add Health and HRS, and are presented for two settings: (1) using all valid samples (“All valid”) and (2) equalizing sample sizes (“Consistent”) across ancestral groups (i.e. randomly selecting a subsample of the European group to match the sample size of the minority populations) in order to ensure that any reduction in prediction accuracy was not due to finite sample bias (Greene 2020).

**Figure 1.**
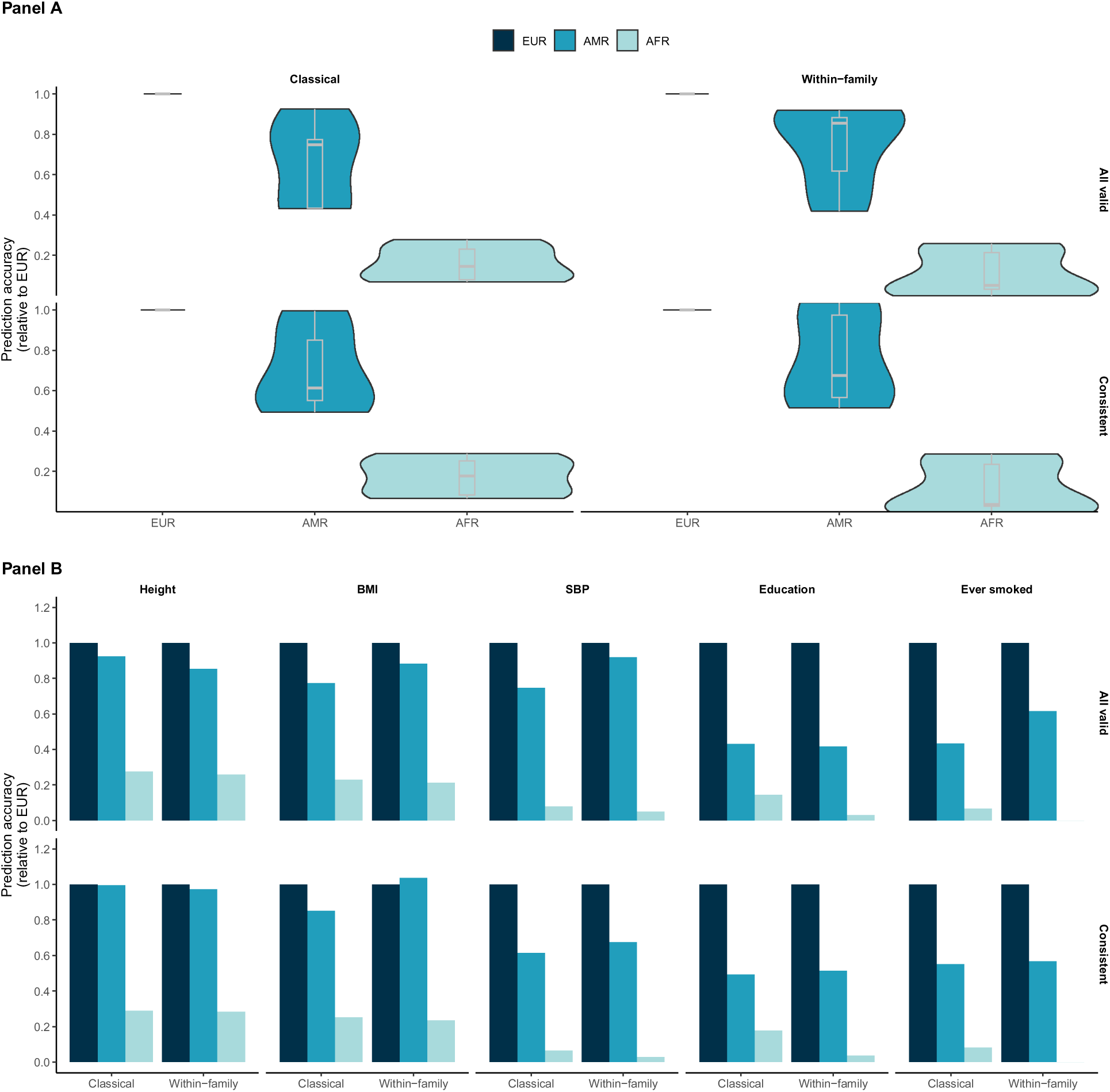
Prediction accuracy of classical and family-based PGIs among the Latino and African American samples relative to the European American sample. Results are meta-analyzed across Add Health and HRS. Panel B presents the relative prediction accuracy for the five phenotypes individually: height, BMI, SBP, educational attainment, and having ever smoked. Panel A displays violin plots summarizing the distribution of relative prediction accuracy across the five phenotypes. The “All valid” rows utilize all available respondents, whereas the “Consistent” rows are based on equalized sample sizes across ancestral groups.

For the Latino group, height shows no diminution. Otherwise, all the phenotypes experience reduced prediction accuracy as measured by the incremental R-squared of the classical PGI when moving from prediction in the European sample to the Latino and African American samples. The decline in R-squared is particularly pronounced for African Americans, as has been demonstrated elsewhere (Martin et al. 2019; Bitarello and Mathieson 2020). When shifting to family-based PGIs, the overall pattern of reduced prediction accuracy persists, with the exception of the BMI PGI, which demonstrates slightly higher accuracy in the Latino group compared to the European group. The upper violin plots in panel A visualize the distribution of the relative prediction accuracy for the five phenotypes. The results clearly indicate a loss of prediction accuracy for both classical and family-based PGIs in the African ancestral group. Importantly, using family-based GWAS weights does not mitigate this decline, underscoring the ongoing challenge of improving PGI portability across diverse ancestries.

Even within genetic ancestral groups, PGI accuracy can vary along the genetic ancestry continuum. Figure 2 illustrates the prediction performance of PGIs for African and Latino American ancestral groups, stratified by their levels of European GSPs. Among both ancestral groups, classical PGIs generally demonstrate higher prediction accuracy in subgroups with greater European GSPs, with statistically significant differences observed for BMI in the African group and for educational attainment and ever smoked status in the Latino group. SBP appears to deviate from this pattern, but further examination with larger sample sizes is needed to ensure sufficient statistical power. Family-based PGIs exhibit similar but slightly less pronounced patterns. However, the BMI PGI still shows significantly better prediction accuracy among African Americans with higher European GSPs.

**Figure 2.**
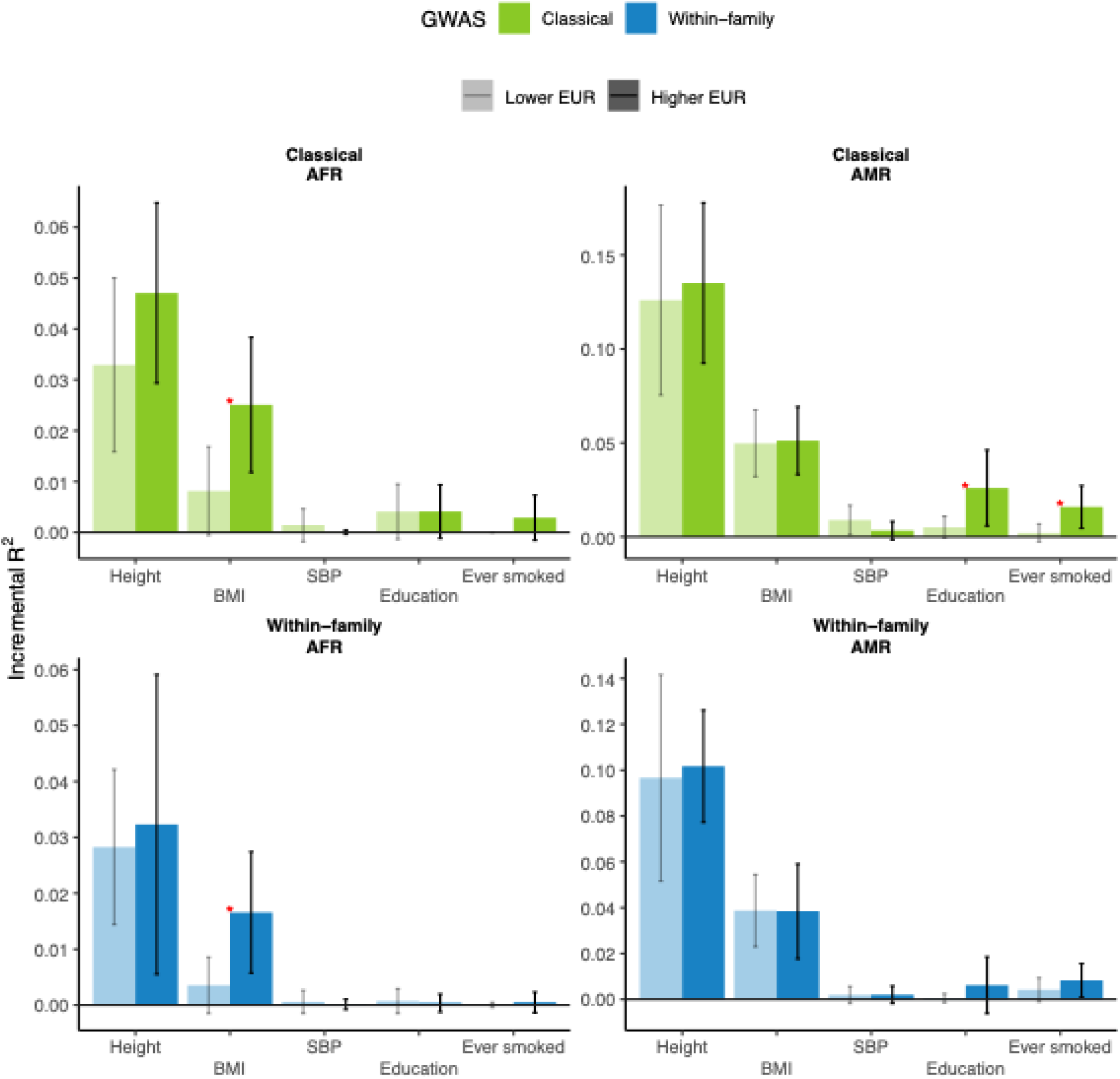
Prediction accuracy of PGIs among African and Latino Americans stratified by European GSPs. The top panels present results for classical PGIs, while the bottom panels show results for family-based PGIs. Left-hand panels pertain to individuals with African genetic ancestry, and right-hand panels to Latino Americans. In each panel, each group is divided into two equal-sized subgroups based on their levels of European GSPs. Differences in prediction accuracy between the subgroups are examined using t-tests. Statistically significant differences at the 0.05 level are marked with red asterisks.

In Figure 3, we present the estimated contributions of variations in genetic architecture, including short-range LD and allele frequencies, to the loss of PGI accuracy in the African ancestral group, following methods developed and detailed in Wang et al (2020). Results for classical GWASs are available for all five phenotypes, while those for within-family GWASs are limited to height and BMI due to the absence of genome-wide significant SNPs for the remaining phenotypes in the study datasets. For height and BMI, more than 85% of the PGI portability issue is attributable to differences in genetic architecture between the European and African genetic ancestral groups. Importantly, the contributions of LD and MAF differences are highly comparable between classical and family-based GWASs for the two phenotypes. For SBP, educational attainment, and having ever smoked, the proportions of reduced accuracy for classical PGIs explained by genetic architecture differences are lower, ranging from 39.8% for educational attainment to 57.4% for having ever smoked. These findings underscore the dominant role of genetic architecture in shaping PGI portability across populations.

**Figure 3.**
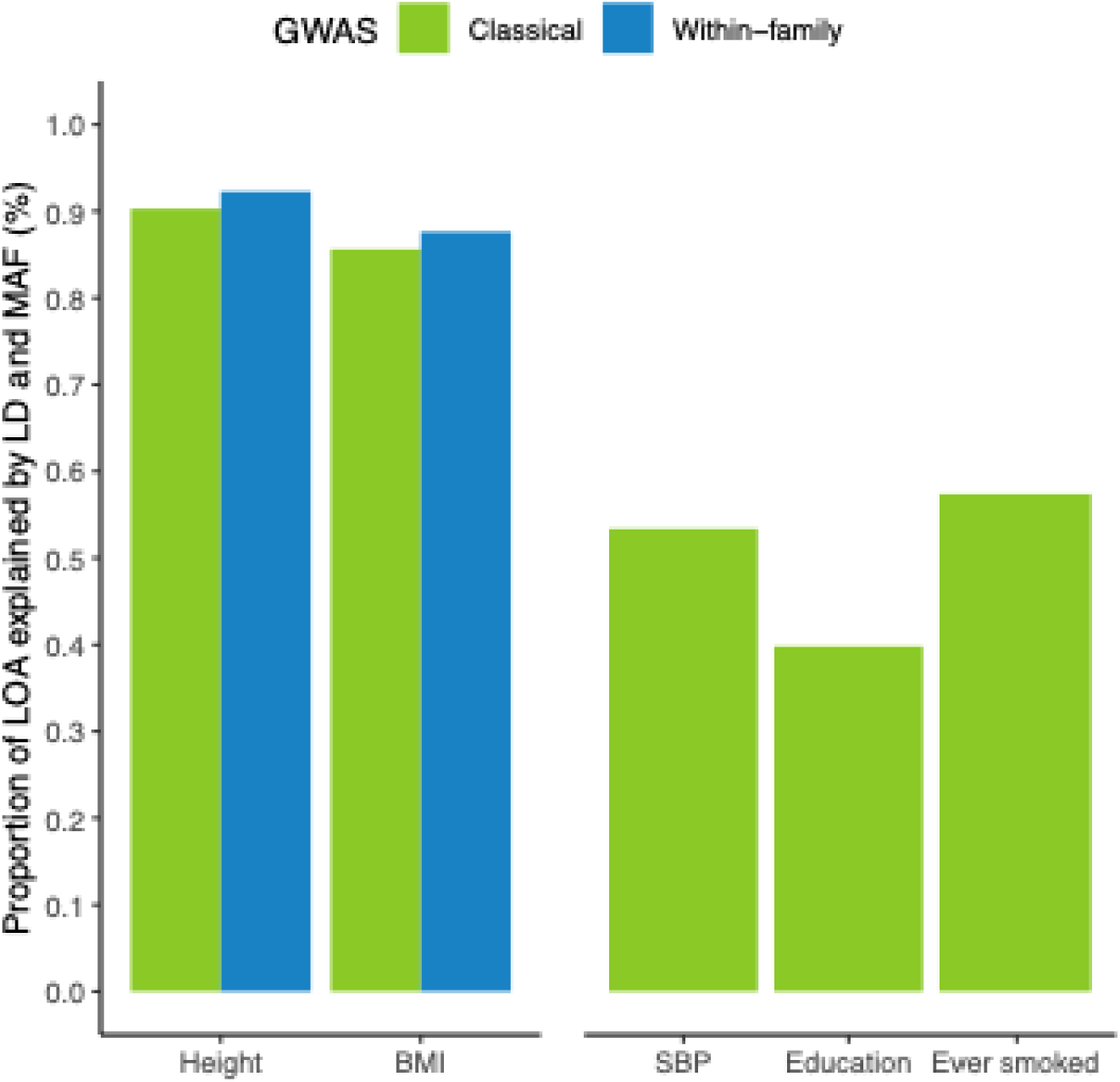
Proportions of loss of prediction accuracy among the African ancestral group explained by LD and MAF differences for the five phenotypes. Calculations follow the methodology in Wang et al. (2020), which considers only genome-wide significant SNPs. Results are provided for classical GWASs across all five phenotypes, while within-family GWASs are limited to height and BMI due to the lack of genome-wide significant SNPs for the other phenotypes in the study datasets.

Finally, in Figure 4, we explore two potential factors influencing the portability of PGIs in the African ancestral group: SNP heritability and assortative mating. Panels A-B reveal that phenotypes with higher heritability in the European and African ancestral groups tend to have PGIs that are more portable to the African sample. Panel C highlights the shrinkage of heritability from classical to family-based GWASs, an indicator of confounding by population stratification, assortative mating, and genetic nurture. Educational attainment shows the largest reduction in heritability, whereas physical traits, including height, BMI, and SBP, experience less diminution. The inverse relationship between heritability shrinkage and PGI portability suggests that lower confounding in classical GWASs is associated with higher portability to the African ancestral group. Panels D-E focus instead on phenotypic assortative mating among the African ancestral group relative to the European ancestral group, measured by sibling correlations in Add Health and spousal correlations in HRS. A higher African-to-European ratio of assortative mating generally corresponds to lower relative PGI accuracy, a pattern consistent across datasets. Notably, this trend holds for classical and family-based PGIs, suggesting that differential assortative mating may not be the primary driver of reduced PGI portability across ancestries.

**Figure 4.**
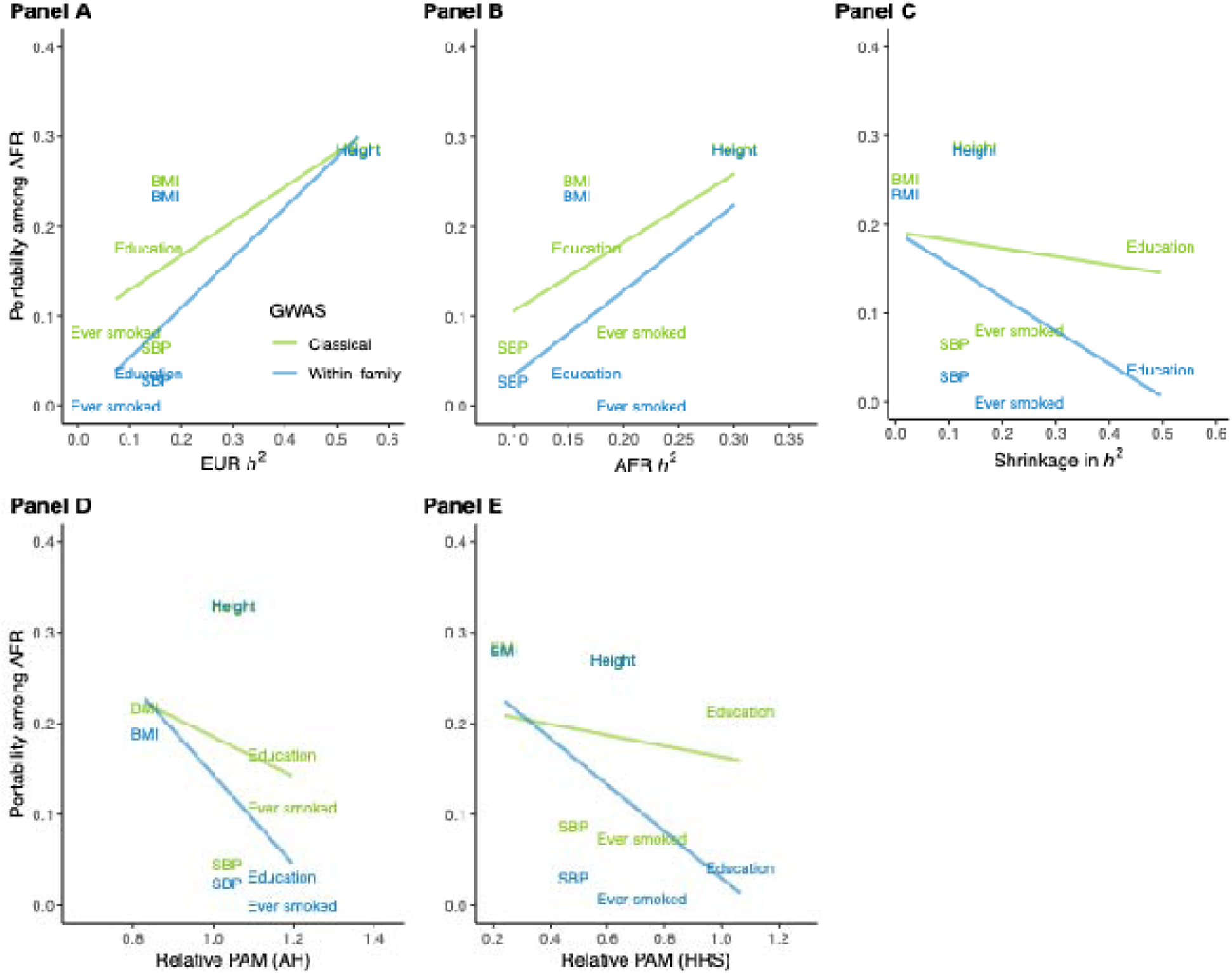
Scatter plots depicting the relative prediction accuracy of PGIs for the African group against measures of heritability and phenotypic assortative mating. Panels A-C visualize relative prediction accuracy against SNP heritability estimates in European and African ancestral groups from the UKB, and the shrinkage in heritability between classical and family-based GWAS (measured as the ratio of classical GWAS heritability to family-based GWAS heritability). The shrinkage is derived from LDSC-based heritability values reported by Tan et al. (2024). Panels D and E display relative prediction accuracy against the AFR-to-EUR ratio of phenotypic assortative mating (PAM) in Add Health (left) and HRS (right), respectively. PAM is measured as sibling correlations in the former and spousal correlations in the latter. Relative prediction accuracy is meta-analyzed across Add Health and HRS for the top three panels.

## Discussion

The lack of portability of polygenic prediction accuracy across groups is of paramount concern as we enter an age where polygenic indices are increasingly used in settings outside the research lab—including, clinical applications, fertility clinics, and social institutions. In this paper we add to the literature examining the root causes of the failure of PGIs trained on samples of exclusively European descent to predict with similar accuracy in other continental ancestry groups (specifically Native Americans as proxied by U.S. Latinos and Africans as represented by ad-mixed African Americans).

The main contribution of this study is testing whether the deployment of PGIs based on within-family GWAS, putatively purged of population stratification, genetic nurture, and assortative mating effects, all of which may bias PGIs differently by population, help ameliorate this decline in prediction accuracy when a PGI is ported across groups. However, within-family-GWAS-based PGIs do no better in stemming the relative decline in prediction accuracy. Additional analysis shows that genetic architecture—rather than GxE or other environmental factors differentially correlated with SNPs across populations—largely accounts for the decline in accuracy. Namely, prediction accuracy generally declines less among those Latinos and African Americans with higher European admixture; this result also obtains in a similar manner whether we use classically based PGIs or within-family PGIs. Such a dynamic, however, might also stem from social factors experienced differently by those with more or less European heritage. Thus, we also directly analyze the contributions of allele frequency differences and differences in LD patterns to the decline in prediction accuracy from European Americans to African Americans. While we could only perform this exercise for two phenotypes – height and BMI – we found that for both the within-family and classically based PGIs the decline in prediction accuracy is largely explained by measures of genetic architecture. Importantly, the differences between the two approaches are trivial. It is worth noting, however, that for the other three phenotypes – SBP, ever smoked and educational attainment – the contribution of MAF and LD is much lower (at least in the classical approach; we could not compute these statistics for the within-family methods given a lack of genome-wide significant SNPs in that analysis).

Finally, among our five phenotypes, we plotted how various other factors affected the degree to which each manifested PGI prediction accuracy declines from the European group to the African American group. There are suggestive patterns. For example, phenotypes with higher heritability tend to port better as do those that experience less shrinkage in heritability moving from classical to within-family GWASs. Meanwhile, those phenotypes with greater assortative mating among African Americans relative to European Americans tend to display greater prediction accuracy decline. However, the most important thing to note in these analyses is that the pattern is the same for the classical PGI as it is for the within-family PGI, suggesting that whatever is driving these patterns is holding true for direct genetic effects.

Purged of population stratification and several other confounders, within-family GWAS is the way of the future for researchers who hope to obtain less biased estimates of SNP or PGI effects. It was with this in mind that we had hypothesized that within-family analysis might also help solve the cross-population portability problem. Alas, whatever their benefits for analyses within a given population, within-family PGIs do not seem to be a panacea for cross-population portability. Of course, it is important to keep in mind two caveats. First, while direct genetic effects estimated among subjects of exclusively European descent are purged of confounding within the sample on which they were estimated, it is possible that these direct effects differently interact with environmental factors in the new population (i.e. Latinos or African Americans).

In this vein, while we compared PGIs weighted by classical versus within-family GWAS in the discovery phase, we did not test within-family approaches in the prediction phase. This is due to the fact that while Add Health had a family structure (specifically a sibling subsample) that would have made such analysis possible, the number of siblings for the Latino and African American populations were prohibitively small for any meaningful comparison (though we show these results in Figure S5 of the Supplementary Material). Future analysis with more data would ideally compare within and between family analysis at both stages—discovery and prediction—to see where prediction accuracy might be affected. This would more directly test whether confounds are introduced at the prediction phase. However, at this point we can be fairly confident in declaring that conducting within-family analysis at the discovery phase does not ameliorate the portability problem.

## Footnotes

5 Height, BMI, systolic blood pressure (SBP), diastolic blood pressure (DBP), cigarettes per day (CPD), number of children ever born (NEB), alcohol consumption, years of schooling, cognitive performance, depressive symptoms, and ever smoked.

6 This is because except for the variants imputed with highest quality, standard imputation from population reference panels do not preserve the relationships between siblings’ genotypes implied by Mendelian Laws.

7 When either of them was missing, the other single measure was used.

8 From Wave 7 (2004) forward, the ever-smoked status was no longer updated for earlier participants, because both the ever and now smoking questions were skipped if the respondent had previously stated having never smoked. However, this will not lead to substantial misclassification, given the observation that respondents who had never smoked in an earlier wave rarely respond “yes” to currently smoking cigarettes in a later wave (RAND 2023).

9 Following Wang et al (2020), African Caribbeans in Barbados (ACB) and Americans of African Ancestry in SW USA (ASW) populations were excluded from AFR.

## Work Cited

Alexander, D.H., Novembre, J. and Lange, K., 2009. Fast model-based estimation of ancestry in unrelated individuals. Genome research, 19(9), pp.1655–1664.

Beauchamp, J.P., 2016. Genetic evidence for natural selection in humans in the contemporary United States. Proceedings of the National Academy of Sciences, 113(28), pp.7774–7779.

Bergström, A., McCarthy, S.A., Hui, R., Almarri, M.A., Ayub, Q., Danecek, P., Chen, Y., Felkel, S., Hallast, P., Kamm, J. and Blanché, H., 2020. Insights into human genetic variation and population history from 929 diverse genomes. Science, 367 (6484), p.eaay5012.

Bitarello, B.D. and Mathieson, I., 2020. Polygenic scores for height in admixed populations. G3: Genes, Genomes, Genetics, 10(11), pp.4027-4036.

Chang, C.C., Chow, C.C., Tellier, L.C., Vattikuti, S., Purcell, S.M. and Lee, J.J., 2015. Second-generation PLINK: rising to the challenge of larger and richer datasets. Gigascience, 4(1), pp.s13742-015.

Ding, Y., Hou, K., Xu, Z., Pimplaskar, A., Petter, E., Boulier, K., Privé, F., Vilhjálmsson, B.J., Olde Loohuis, L.M. and Pasaniuc, B., 2023. Polygenic scoring accuracy varies across the genetic ancestry continuum. Nature, 618(7966), pp.774–781.

Ge, T., Chen, C.Y., Ni, Y., Feng, Y.C.A. and Smoller, J.W., 2019. Polygenic prediction via Bayesian regression and continuous shrinkage priors. Nature communications, 10(1), p.1776.

Greene, W.H. 2020. Econometric Analysis. 8th ed. New York: Pearson Education.

Health and Retirement Study, (RAND HRS Longitudinal File 2020 (V1)) public use dataset. Produced and distributed by the University of Michigan with funding from the National Institute on Aging (grant number NIA U01AG009740). Ann Arbor, MI, (March 2023).

Hilmarsson, H., Kumar, A.S., Rastogi, R., Bustamante, C.D., Montserrat, D.M. and Ioannidis, A.G., 2021. High resolution ancestry deconvolution for next generation genomic data. bioRxiv, pp.2021–09.

Hinrichs, A.S., Karolchik, D., Baertsch, R., Barber, G.P., Bejerano, G., Clawson, H., Diekhans, M., Furey, T.S., Harte, R.A., Hsu, F. and Hillman-Jackson, J., 2006. The UCSC genome browser database: update 2006. Nucleic acids research, 34(suppl_1), pp.D590-D598.

Howe, L.J., Nivard, M.G., Morris, T.T., Hansen, A.F., Rasheed, H., Cho, Y., Chittoor, G., Ahlskog, R., Lind, P.A., Palviainen, T. and van der Zee, M.D., 2022. Within-sibship genome-wide association analyses decrease bias in estimates of direct genetic effects. Nature genetics, 54(5), pp.581–592.

Howie, B.N., Donnelly, P. and Marchini, J., 2009. A flexible and accurate genotype imputation method for the next generation of genome-wide association studies. PLoS genetics, 5(6), p.e1000529.

International HapMap 3 Consortium, 2010. Integrating common and rare genetic variation in diverse human populations. Nature, 467 (7311), p.52.

Kumar, A., Im, K., Banjevic, M., Ng, P.C., Tunstall, T., Garcia, G., Galhardo, L., Sun, J., Schaedel, O.N., Levy, B. and Hongo, D., 2022. Whole-genome risk prediction of common diseases in human preimplantation embryos. Nature medicine, 28 (3), pp.513–516.

Manichaikul, A., Mychaleckyj, J.C., Rich, S.S., Daly, K., Sale, M. and Chen, W.M., 2010. Robust relationship inference in genome-wide association studies. Bioinformatics, 26(22), pp.2867–2873.

Martin, A.R., Gignoux, C.R., Walters, R.K., Wojcik, G.L., Neale, B.M., Gravel, S., Daly, M.J., Bustamante, C.D. and Kenny, E.E., 2017. Human demographic history impacts genetic risk prediction across diverse populations. The American Journal of Human Genetics, 100 (4), pp.635–649.

Martin, A.R., Kanai, M., Kamatani, Y., Okada, Y., Neale, B.M. and Daly, M.J., 2019. Clinical use of current polygenic risk scores may exacerbate health disparities. Nature genetics, 51(4), pp.584–591.

Monk Jr, E.P., 2021. The unceasing significance of colorism: Skin tone stratification in the United States. Daedalus, 150(2), pp.76–90.

Mostafavi, H., Harpak, A., Agarwal, I., Conley, D., Pritchard, J.K. and Przeworski, M., 2020. Variable prediction accuracy of polygenic scores within an ancestry group. elife, 9, p.e48376.

O’Sullivan, J.W., Raghavan, S., Marquez-Luna, C., Luzum, J.A., Damrauer, S.M., Ashley, E.A., O’Donnell, C.J., Willer, C.J. and Natarajan, P., 2022. Polygenic risk scores for cardiovascular disease: a scientific statement from the American Heart Association. Circulation, 146(8), pp.e93–e118.

Peterson, R.E., Kuchenbaecker, K., Walters, R.K., Chen, C.Y., Popejoy, A.B., Periyasamy, S., Lam, M., Iyegbe, C., Strawbridge, R.J., Brick, L. and Carey, C.E., 2019. Genome-wide association studies in ancestrally diverse populations: opportunities, methods, pitfalls, and recommendations. Cell, 179(3), pp.589–603.

Purcell, S., Neale, B., Todd-Brown, K., Thomas, L., Ferreira, M.A., Bender, D., Maller, J., Sklar, P., De Bakker, P.I., Daly, M.J. and Sham, P.C., 2007. PLINK: a tool set for whole-genome association and population-based linkage analyses. The American journal of human genetics, 81(3), pp.559–575.

RAND HRS Longitudinal File 2020 (V1). Produced by the RAND Center for the Study of Aging, with funding from the National Institute on Aging and the Social Security Administration. Santa Monica, CA (March 2023).

Ruderfer, D.M., Ripke, S., McQuillin, A., Boocock, J., Stahl, E.A., Pavlides, J.M.W., Mullins, N., Charney, A.W., Ori, A.P., Loohuis, L.M.O. and Domenici, E., 2018. Genomic dissection of bipolar disorder and schizophrenia, including 28 subphenotypes. Cell, 173(7), pp.1705–1715.

Tan, T., Jayashankar, H., Guan, J., Nehzati, S.M., Mir, M., Bennett, M., Agerbo, E., Ahlskog, R., Pinto de Andrade Anapaz, V., Asvold, B.O. and Benonisdottir, S., 2024. Family-GWAS reveals effects of environment and mating on genetic associations. medRxiv, pp.2024–10.

The International Schizophrenia Consortium. Common polygenic variation contributes to risk of schizophrenia and bipolar disorder. Nature 460, 748–752 (2009). 10.1038/nature08185

Turley, P., Meyer, M.N., Wang, N., Cesarini, D., Hammonds, E., Martin, A.R., Neale, B.M., Rehm, H.L., Wilkins-Haug, L., Benjamin, D.J. and Hyman, S., 2021. Problems with using polygenic scores to select embryos. New England Journal of Medicine, 385 (1), pp.78–86.

United Nations, Department of Economic and Social Affairs, Population Division. (2024). World Population Prospects 2024: Summary of Results. Available at: https://population.un.org/wpp/Publications/Files/WPP2024_Summary_of_Results.pdf [Accessed 19 Nov. 2024].

Wang, Y., Guo, J., Ni, G., Yang, J., Visscher, P.M. and Yengo, L., 2020. Theoretical and empirical quantification of the accuracy of polygenic scores in ancestry divergent populations. Nature communications, 11(1), p.3865.

Yang, J., Lee, S.H., Goddard, M.E. and Visscher, P.M., 2011. GCTA: a tool for genome-wide complex trait analysis. The American Journal of Human Genetics, 88(1), pp.76–82.

Yengo, L., Sidorenko, J., Kemper, K.E., Zheng, Z., Wood, A.R., Weedon, M.N., Frayling, T.M., Hirschhorn, J., Yang, J., Visscher, P.M. and Giant Consortium, 2018. Meta-analysis of genome-wide association studies for height and body mass index in∼ 700000 individuals of European ancestry. Human molecular genetics, 27(20), pp.3641–3649.

Young, A.I., Nehzati, S.M., Benonisdottir, S., Okbay, A., Jayashankar, H., Lee, C., Cesarini, D., Benjamin, D.J., Turley, P. and Kong, A., 2022. Mendelian imputation of parental genotypes improves estimates of direct genetic effects. Nature genetics, 54(6), pp.897–905.

1000 Genomes Project Consortium, 2015. A global reference for human genetic variation. Nature, 526 (7571), p.68.

