## Supplemental Figures and Tables for "Within-Family GWAS does not Ameliorate the Decline in Prediction Accuracy across Populations"

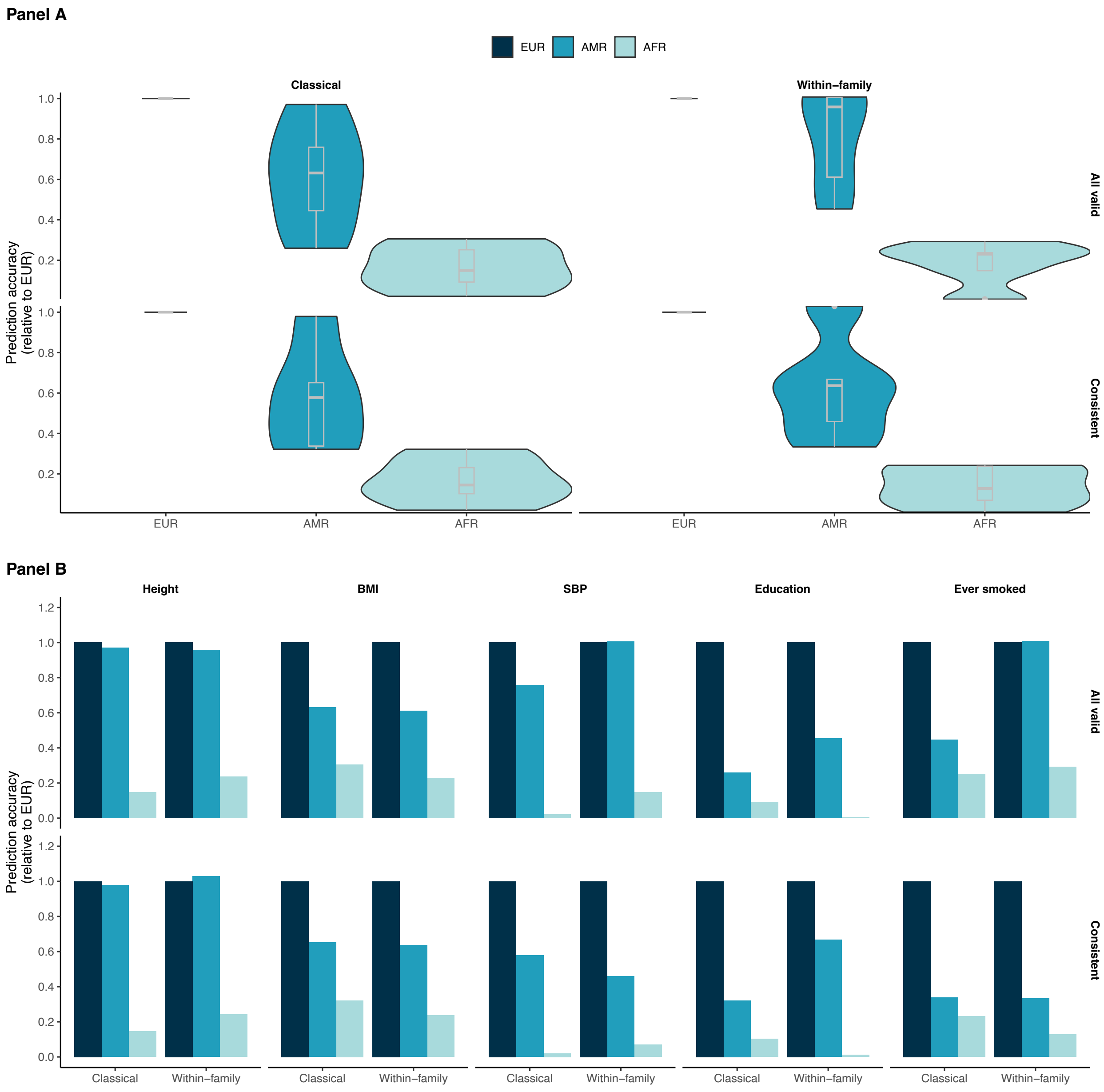


**Figure S1. Relative prediction accuracy of classical and family-based PGIs among the Latino and African American samples based on GWAS summary statistics by Howe et al (2022).** Results are meta-analyzed across Add Health and HRS. Panel A displays violin plots summarizing the distribution of relative prediction accuracy across the five phenotypes. Panel B presents the relative prediction accuracy for the five phenotypes individually: height, BMI, SBP, educational attainment, and having ever smoked. The “All valid” rows utilize all available respondents, whereas the “Consistent” rows are based on equalized sample sizes across ancestral groups.

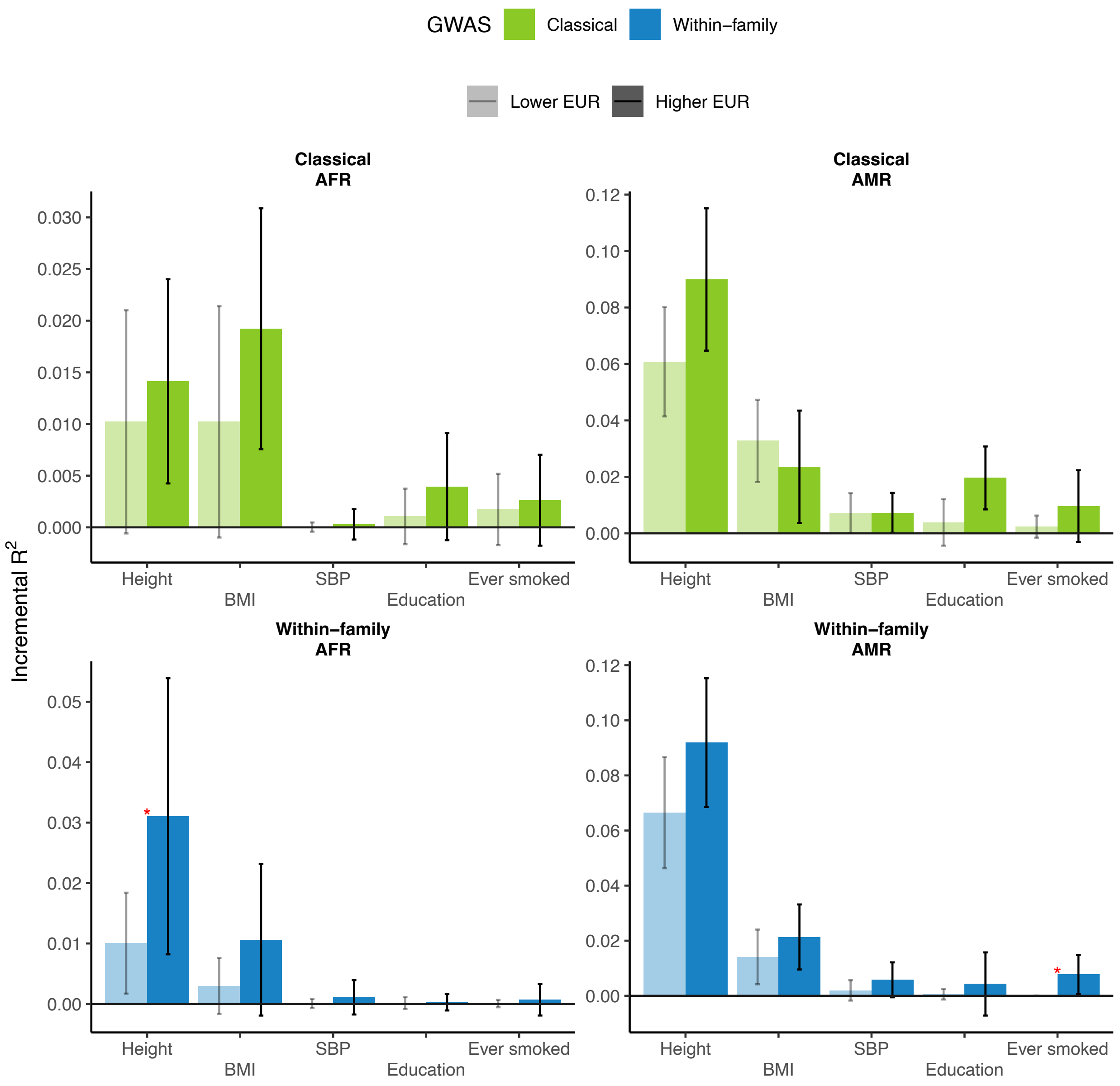


**Figure S2. Prediction accuracy of Howe-based PGIs among African and Latino Americans stratified by European GSPs**. The top panels present results for classical PGIs, while the bottom panels show results for family-based PGIs. Left-hand panels pertain to individuals with African genetic ancestry, and right-hand panels to Latino Americans. In each panel, each group is divided into two equal-sized subgroups based on their levels of European GSPs. Differences in prediction accuracy between the subgroups are examined using t-tests. Statistically significant differences at the 0.05 level are marked with red asterisks.

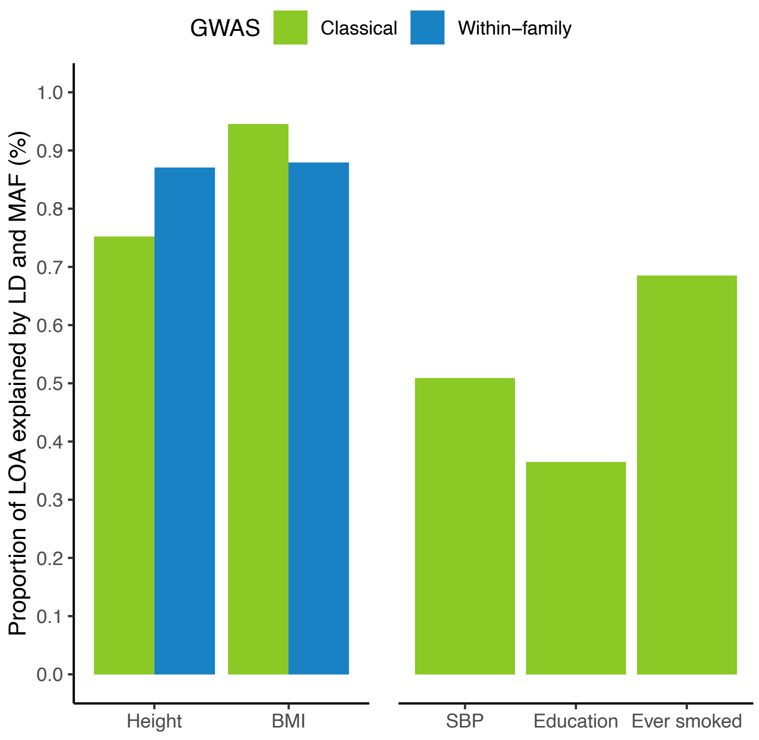


**Figure S3. Proportions of loss of prediction accuracy of Howe-based PGIs among the African ancestral group explained by LD and MAF differences for the five phenotypes**. Calculations follow the methodology in Wang et al. (2020), which considers only genome-wide significant SNPs. Results are provided for classical GWASs across all five phenotypes, while within-family GWASs are limited to height and BMI due to the lack of genome-wide significant SNPs for the other phenotypes in the study datasets.

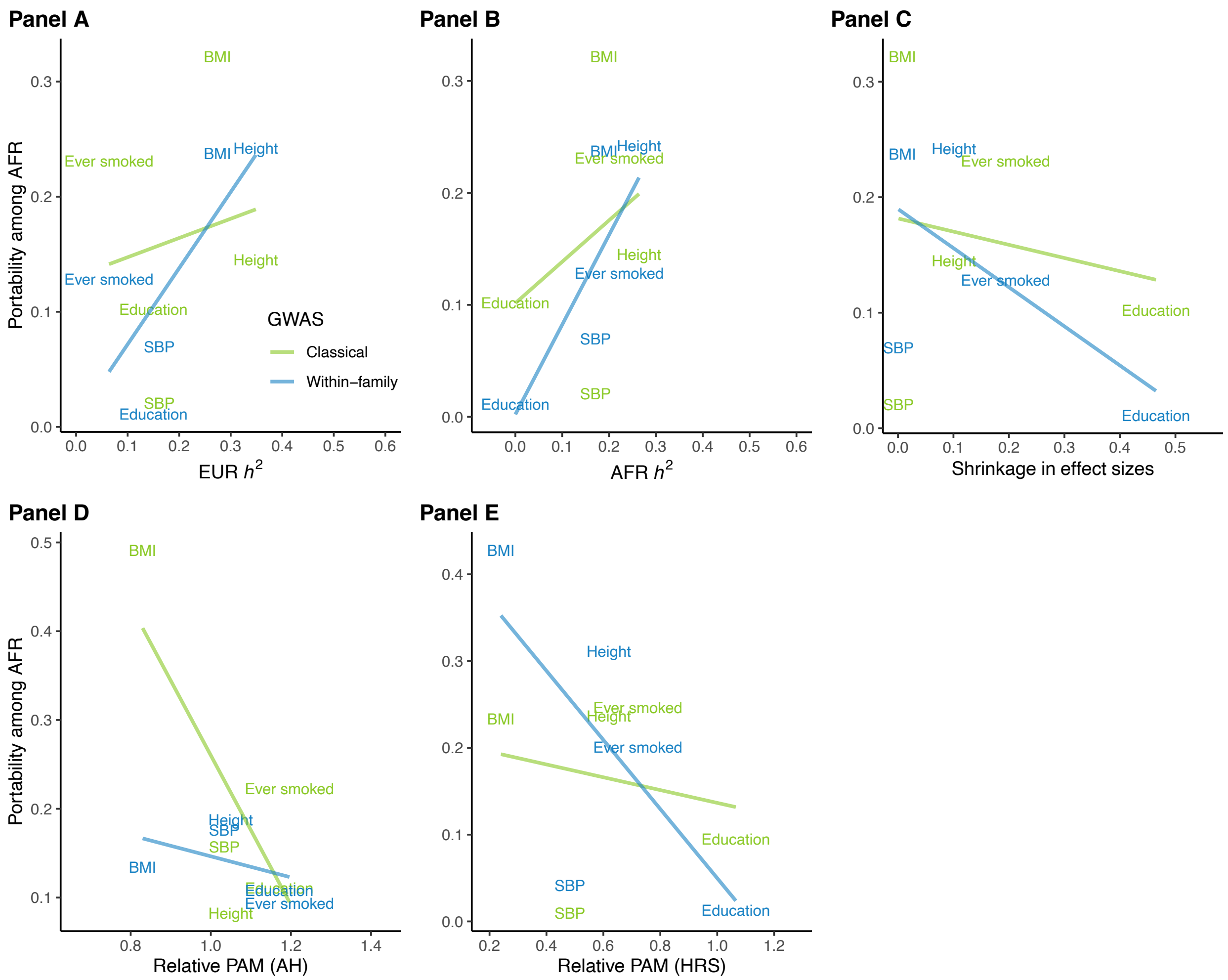


**Figure S4. Scatter plots depicting the relative prediction accuracy of Howe-based PGIs for the African group against measures of heritability and phenotypic assortative mating.** Panels A-B visualize relative prediction accuracy against SNP heritability estimates in European and African ancestral groups from the UKB. Panel C presents relative prediction accuracy against the shrinkage in effect sizes between classical and within-sibship GWASs, which is obtained directly from Howe et al. (2022). Panels D and E display relative prediction accuracy against the AFR-to-EUR ratio of phenotypic assortative mating (PAM) in Add Health (left) and HRS (right), respectively. PAM is measured as sibling correlations in the former and spousal correlations in the latter. Relative prediction accuracy is meta-analyzed across Add Health and HRS for the top three panels.

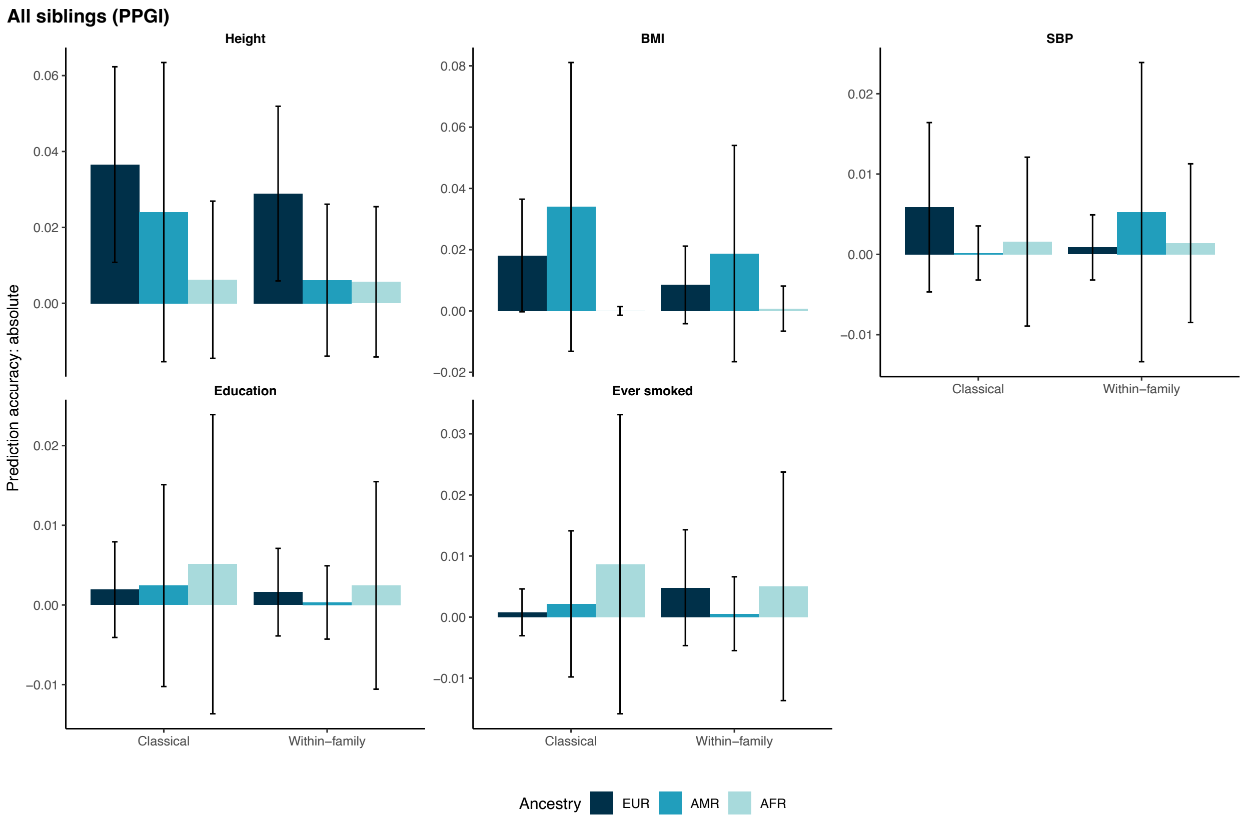


**Figure S5. Prediction accuracy of classical and family-based PGIs in within-family analysis of Add Health siblings**. Results are shown for the five phenotypes (height, BMI, SBP, educational attainment, and having ever smoked) and the three genetic ancestral groups (European, African, and Latino Americans). Prediction accuracy is measured as squared partial correlation between the PGI and the corresponding phenotype after they have been both residualized using gender, birth year, the first 10 genetic PCs, and the imputed paternal PGI.

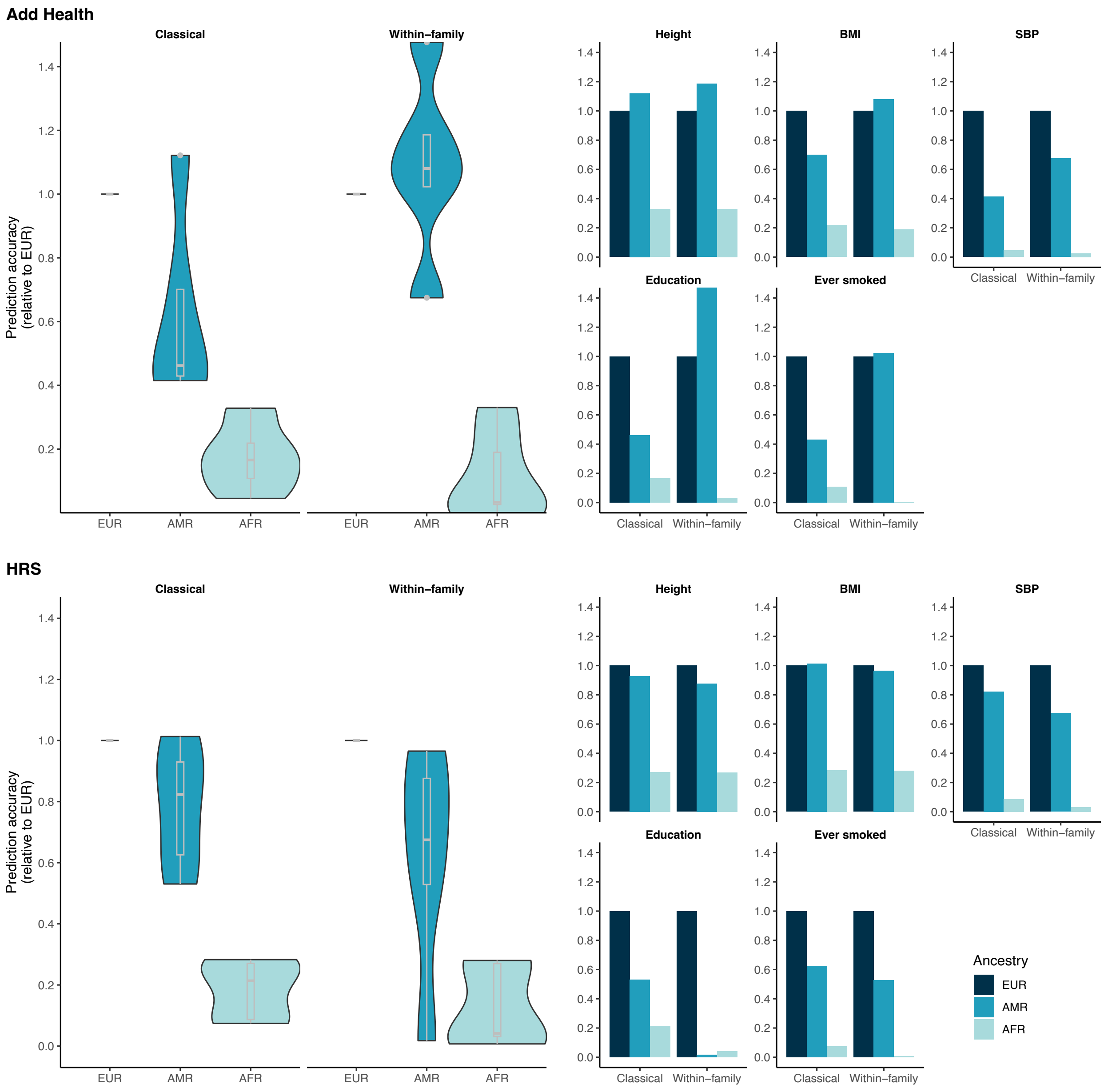


**Figure S6. Relative prediction accuracy of classical and family-based PGIs among the Latino and African American samples in Add Health and HRS**. Results are separately displayed for Add Health and HRS. Right panels present the relative prediction accuracy for the five phenotypes individually: height, BMI, SBP, educational attainment, and having ever smoked. Left panels display violin plots summarizing the distribution of relative prediction accuracy across the five phenotypes. Sample sizes are equalized across genetic ancestral groups.

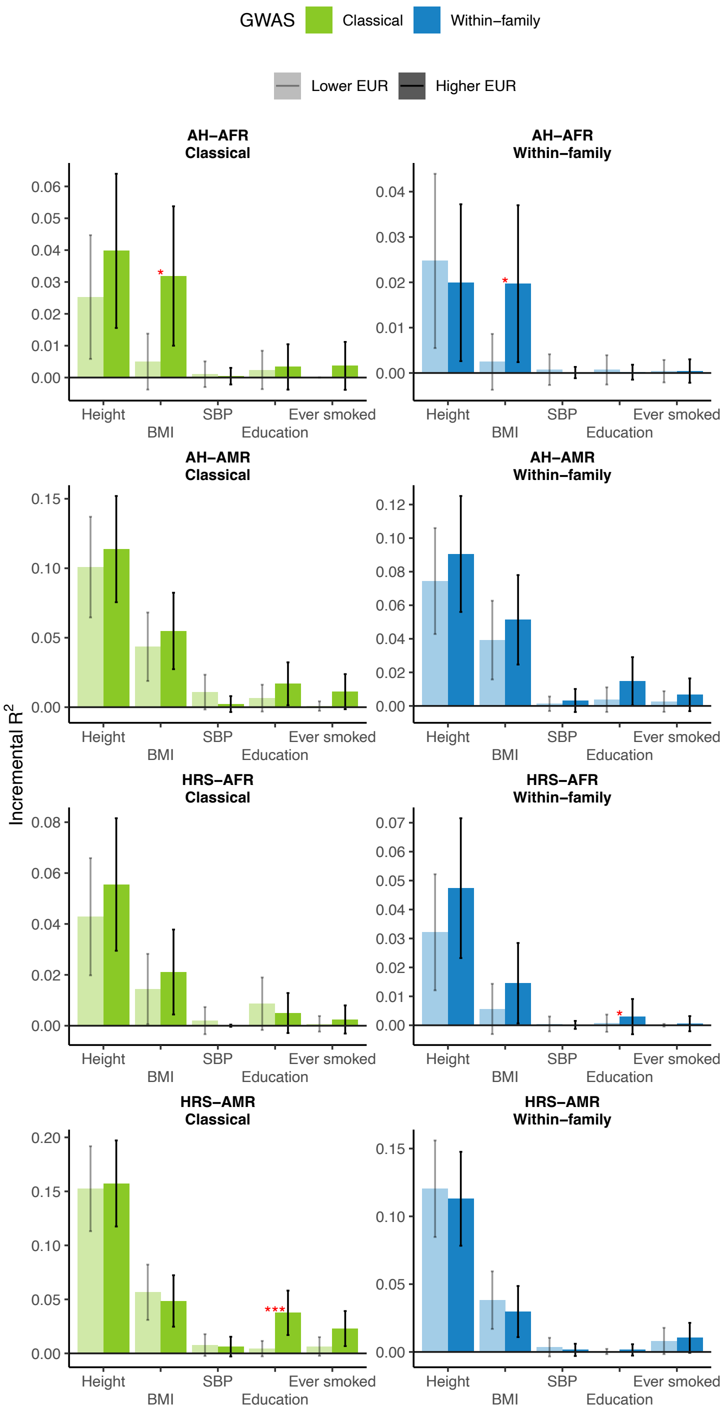


**Figure S7. Prediction accuracy of PGIs among African and Latino Americans in Add Health and HRS stratified by European GSPs**. The top four panels present results from Add Health, while the bottom four panels are based on HRS. Classical and family-based PGIs are shown in green and blue, respectively. In each panel, the respective ancestral group is divided into two equal-sized subgroups based on their levels of European GSPs. Differences in prediction accuracy between the subgroups are examined using t-tests. Statistically significant differences at the 0.05 level are marked with red asterisks.

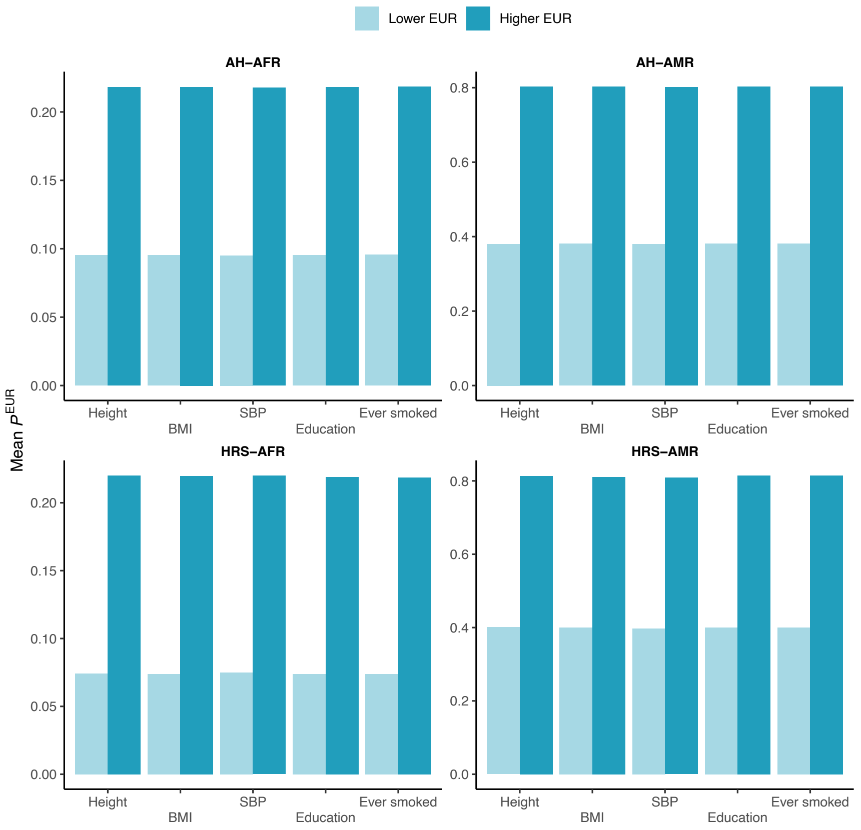


**Figure S8. Average European GSPs of the two equal-sized subgroups of African and Latino Americans in Add Health and HRS**. The top two panels are based on Add Health, while the bottom two panels pertain to HRS respondents. In each panel, mean European GSPs are displayed for the analytical sample of each of the five phenotypes: height, BMI, SBP, educational attainment, and having ever smoked.

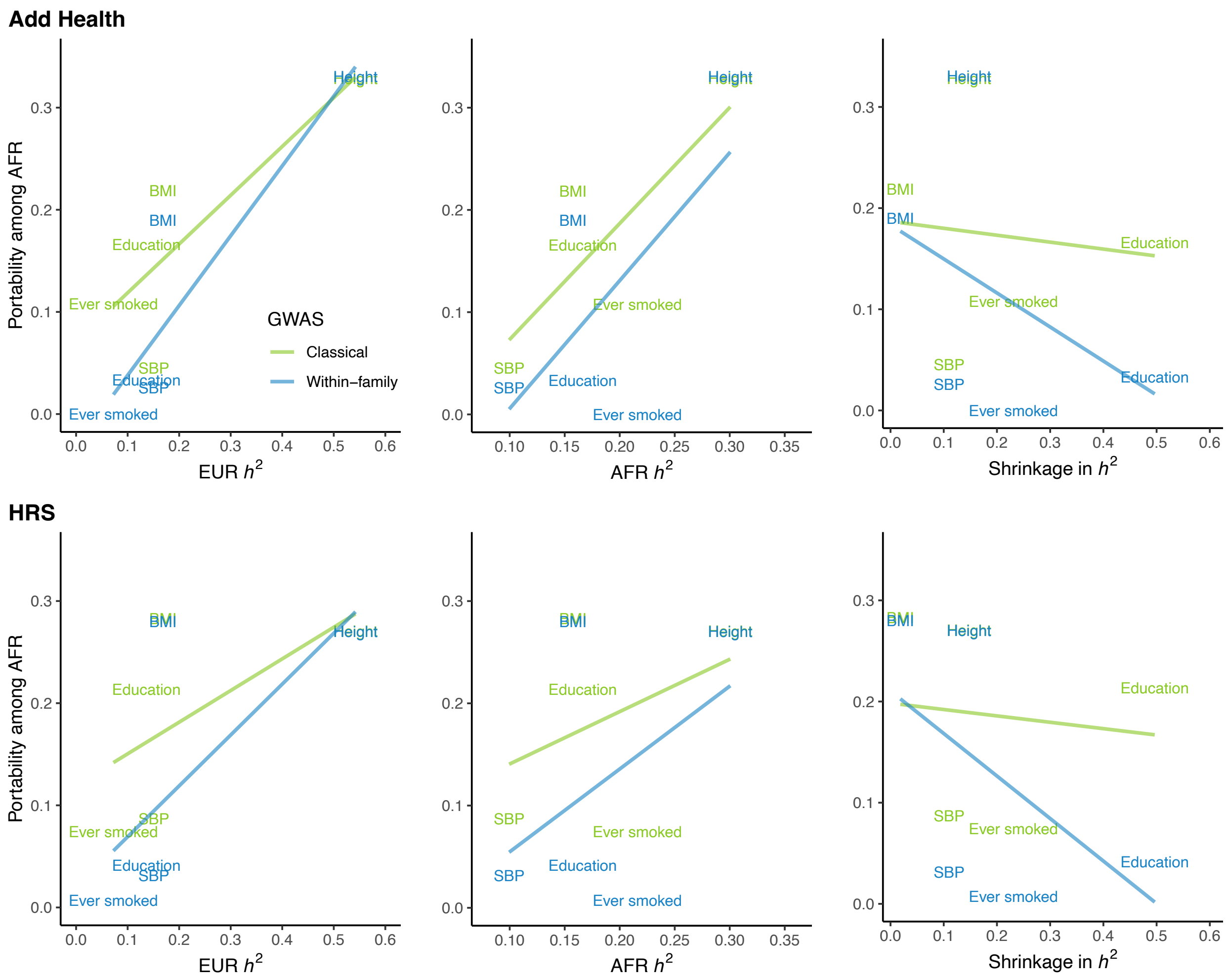


**Figure S9. Scatter plots depicting the relative prediction accuracy of PGIs for the African group in Add Health and HRS against measures of heritability**. The top row shows results from Add Health, while the bottom row presents patterns among HRS respondents. For each dataset, relative prediction accuracy is plotted against SNP heritability estimates in European and African ancestral groups from the UKB, and the shrinkage in heritability between classical and family-based GWASs, which is derived from LDSC-based heritability values reported by Tan et al. (2024).
